# Dehydration alters transcript levels in the mosquito midgut, likely facilitating rapid rehydration

**DOI:** 10.1101/2022.09.15.508146

**Authors:** Christopher J. Holmes, Elliott S. Brown, Dhriti Sharma, Matthew Warden, Atit Pathak, Blaine Payton, Quynh Nguyen, Austin Spangler, Jaishna Sivakumar, Jacob M. Hendershot, Joshua B. Benoit

## Abstract

The mosquito midgut is an important site for bloodmeal regulation while also acting as a primary site for pathogen exposure within the mosquito. Recent studies show that exposure to dehydrating conditions alters mosquito bloodfeeding behaviors as well as post-feeding regulation, likely altering how pathogens interact with the mosquito. Unfortunately, few studies have explored the underlying dynamics between dehydration and bloodmeal utilization, and the overall impact on disease transmission dynamics remains veiled. In this study, we find that dehydration-based feeding in the yellow fever mosquito, *Aedes aegypti*, prompts alterations to midgut gene expression, as well as subsequent physiological factors involving water control and post-bloodfeeding (pbf) regulation. Altered expression of ion transporter genes in the midgut of dehydrated mosquitoes and rapid reequilibration of hemolymph osmolality after a bloodmeal indicate an ability to expedite fluid and ion processing. These alterations ultimately indicate that female *A. aegypti* employ mechanisms to ameliorate the detriments of dehydration by imbibing a bloodmeal, providing an effective avenue for rehydration. Continued research into bloodmeal utilization and the resulting effects on arthropod-borne transmission dynamics becomes increasingly important as drought prevalence is increased by climate change.

## Introduction

Numerous studies over the last century have investigated the relationships between mosquitoes and relative humidity [1–15]. However, only a subset of those studies has investigated the physiological effects of low relative humidity on mosquito biology, with an even smaller subset controlling for and directly studying the impacts of relative humidity on mosquitoes. This disparity warrants further exploration, especially considering that weather conditions are a direct cause of dehydration in mosquitoes, and that incorporation of weather conditions into models may account for up to 80% of the weekly variation in mosquito infection [1,16].

Recent studies implicate dehydration stress in water and nutrient depletion, as well as in the compensatory mechanisms (e.g., increased bloodmeal retention) required to offset those detriments [1,7,17]. Unfortunately, these identified mechanisms have been predicted to alter disease propagation dynamics both within the vector and through host-vector interactions [1,7]. For example, previous findings indicate that nutrient reserves in the northern house mosquito, *Culex pipiens*, decreased as dehydration exposure increased, resulting in reductions to mosquito survival and reproduction [8]. Conversely, fortified nutritional reserves have been shown to improve longevity and increase resistance to pathogen challenge [18]; but direct connections between dehydration and disease transmission dynamics remains unexplored. It is therefore paramount to understand the specifics on how humidity drives alterations in mosquito physiology as well as the biological components and underlying compensatory mechanisms required to offset any related detriments.

Compensatory behaviors are well documented within mosquitoes, with an early study on *Anopheles* species showing that blood digestion increased during the hot season [15] and later studies demonstrating that a bloodmeal could be utilized for nutritional supplementation [19,20]. Hagan et al. (2018) began investigating the potential for compensatory mechanisms in dehydrated mosquitoes, finding that biting propensity and carbohydrate metabolism was altered in dehydrated *C. pipiens*, culminating in a predicted increase to West Nile virus (WNV) transmission [1]. Holmes et al. (2022) continued this line of research, finding in a recent study with *C. pipiens* and *A. aegypti* that dehydration prompted increases in bloodfeeding propensity and greater water content retention from a bloodmeal, resulting in improved survival for bloodfed mosquitoes in dehydrating conditions [7]. These responses to dehydration were predicted to increase compensatory bloodfeeding as a response to lost water, ultimately altering the vectorial capacity of both *C. pipiens* and *A. aegypti* [7].

When incorporated into disease models, transmission has been found to be strongly influenced, and predicted, by factors such as environmental stressors [21], viral transmission [22,23], differential expression of genes [24], and the interactions between those factors [1]. Considering the reliance of various disease transmission models on relative humidity as a factor, as well as the numerous implications of relative humidity on mosquito physiology and behavior [17], more research must be aimed at addressing the direct effects of water loss (i.e., dehydration) on mosquitoes. To continue addressing this lapse in research, our study incorporated transcriptomic analyses and physiological assays to address the biological effects of dehydration stress on *A. aegypti* bloodmeal processing. Specifically, this study developed transcriptomic profiles for the midguts of *A. aegypti* subjected to dehydration stress in relation to bloodfeeding, facilitating a better understanding of the compensatory mechanisms underlying physiological alterations. Understanding the interactions of a bloodmeal within the midgut of a dehydrated mosquito may offer insights into potential permissibility differences in the gut (e.g., through altered regulatory mechanisms), with possible implications for disease transmission dynamics. Regardless, understanding the effect that a natural stressor like dehydration has on the midgut further necessitates the inclusion of environmental effects in disease dynamics. This study used next-generation sequencing to determine underlying genes involved in post-dehydration bloodmeal regulation in *A. aegypti*. The results of this experiment revealed ion transporters, RNA regulation, and kinase involvement in dehydration and bloodfeeding exposures within the midgut. These findings, in addition those of stabilizing osmolality and unaltered midgut size or micronutrients, provide a more thorough understanding of the mechanisms that drive fluid acquisition and retention in dehydrated mosquitoes.

## Materials and Methods

### Mosquito husbandry

Mosquito larvae were reared according to standard practices on ground fish food (Tetramin) with added yeast extract (Fisher). Adult *A. aegypti* mosquitoes (Rockefeller strain) were reared under insectary conditions (27°C, 80% RH; saturation vapor pressure deficit (SVPD) = 0.71 kPa) in 12 × 12 × 12” cages (BioQuip) with a 16h:8h light:dark cycle and unlimited access to DI water- and 10% sucrose solution-soaked cotton wicks *ad libitum*, unless otherwise stated.

### Relative humidity exposure protocol

Similar to Holmes et al, (2022), mosquitoes were subjected to desiccators containing controlled relative humidity conditions at 27°C with 75% RH (dehydrating condition; SVPD = 0.89 kPa) or 100% RH (non-dehydrating condition; SVPD = 0.00 kPa) by being placed in groups of 50 into mesh-covered 50mL centrifuge tubes. These humidity-controlled mosquitoes were held under desiccator conditions without access to water or sucrose solution for 18 hours before being subjected to downstream procedures.

### Mosquito midgut processing for transcriptomic analyses

After RH treatment, mosquitoes were released into 12 × 12 × 12” cages (BioQuip) and permitted to bloodfeed to repletion (approximately 20 minutes) on a live human host (27-year-old male, leg; IRB, University of Cincinnati) or not permitted to bloodfeed but with a human leg just outside the cage. These conditions resulted in four different groups: N1, non-bloodfed/non-dehydrated (control) group; Y1, bloodfed/non-dehydrated group; N7, non-bloodfed/dehydrated group; Y7, bloodfed/dehydrated group. Three hours (±1h) pbf, mosquitoes were dissected and the midguts from approximately 15 different mosquitoes were pooled and placed into Trizol (Invitrogen) held on ice. Digestion of blood occurs around 4 hours pbf [25] and diuresis is well underway within 2h [26,27], so dissections 3h post-bloodmeal were chosen to encompass differentially expressed genes related to altered blood digestion/water retention. Pooled midguts were homogenized (Benchmark, BeadBlaster 24), in Trizol and stored at −70°C until all samples were collected. RNA was extracted with Trizol according to manufacturer’s protocols and cleaned with a RNeasy Mini Kit (Qiagen). DNase (Ambion, Turbo-DNA-free) was used to remove genomic DNA, RNA concentration was determined with a Nanodrop 2000 (Fisher), cDNA libraries were generated (Illumina, TruSeq), and next-generation sequencing was conducted at the Cincinnati Children’s Hospital Medical Center’s DNA Sequencing and Genotyping Core. Samples can be found in the Sequence Read Archive (SRA) Database (BioProject ID: PRJNA851095).

### Gene expression analyses

Samples were analyzed through three separate pipelines using recommended settings throughout: CLC Genomics Workbench 12.1 (CLC Bio, Boston, MA, USA), DESeq2-Kallisto, and DESeq2-Sailfish. All pipelines used the published *A. aegypti* RefSeq assembly (accession: GCF_002204515.2) as reference [28]. The latter two pipelines included importing samples into Galaxy [29], checking for quality with FastQC [30], trimming with Trimmomatic [31], and analyzing with Kallisto [32] or Sailfish [33], before utilization of DESeq2 [34]. Significantly expressed genes were determined by Bonferroni correction (p-value < 0.01), the genes identified by any pipeline are provided in (Supplementary Table 1), and the DESeq2 pipeline comparisons between transcript mean expression and fold-changes are included in (Supplementary Table 2). Transcriptomic methods revealed sufficient coverage, with approximately 75-105 million paired-end reads per sample (Table 1). Gene ontology (GO) terms were generated by importing all significantly expressed genes (p-value < 0.01) with a ≥ 2-fold fold-change identified by any pipeline (Supplementary Table 3) into g:Profiler [35]. Gene ontology terms were subsequently summarized with REVIGO [36] and visualized via CirGO [37] (Supplementary Table 4). Although all pipelines were used to identify genes for the GO analyses, only DESeq2 pipeline results were compared for downstream expressional analyses. The CLC pipeline protocol included calculated mean expression values of zero for numerous genes, resulting in comparative fold-changes of infinity. However, in the DESeq2 pipelines, genes with expression values of zero were not included as part of the analysis, reducing the false positive identification rate of differentially expressed genes. Due to our smaller sample sizes and these differences in pipeline methodology, only the more conservative DESeq2 pipelines were utilized for further analysis. All log_2_ normalized mean expression values, regardless of group comparison, were compared between the DESeq2-Kallisto and DESeq2-Sailfish pipelines and were found to be considerably correlated (n = 181, r = 0.921, p-value < 0.00001; Supplementary Figure 1).

**Table 1:** Descriptive information regarding sample composition and read counts of experimental groups. Sample numbers are provided in the respective column.

| Group | Dehydration | Bloodfed | Sample | Paired-End Reads |
| --- | --- | --- | --- | --- |
| N1 | No | No | N1-2 | 75,496,800 |
|  |  |  | N1-3 | 88,761,156 |
| Y1 | No | Yes | Y1-1 | 89,322,470 |
|  |  |  | Y1-2 | 81,691,584 |
|  |  |  | Y1-3 | 74,120,060 |
| N7 | Yes | No | N7-1 | 105,818,594 |
|  |  |  | N7-2 | 105,385,942 |
|  |  |  | N7-3 | 82,647,520 |
| Y7 | Yes | Yes | Y7-1 | 95,531,032 |
|  |  |  | Y7-2 | 85,314,086 |

### Osmolality procedures

In addition to the two RH treatments, an additional post-dehydration exposure group was also analyzed 1h after taking a bloodmeal. Bloodfeeding was completed by filling artificial (Hemotek) reservoirs with chicken blood (Pel-Freez Biologicals), covering with parafilm (Sigma-Aldrich), warming to 37°C, introducing the covered reservoir to 12 × 12 × 12” cages (BioQuip) without access to water or sucrose solution for 1h, and allowing the dehydrated mosquitoes to feed to repletion [38]. Before use, chicken blood was held at −20°C and then permitted to thaw at 4°C. One hour after conclusion of RH treatment or post-RH treatment blood feeding, mosquito hemolymph was extracted for osmolality measurement with a vapor pressure osmometer (Wescor Vapro 5600, EliTech).

### Midgut volume quantification

Mosquitoes were bloodfed as before with an artificial feeder (Hemotek) filled with chicken blood (Pel-Freez Biologicals). Within 1h pbf, mosquitoes were knocked out with CO_2_, dissected (N = 86) in phosphate-buffered saline (PBS), and photographed (Dino-Lite). Micrometer measurements were calibrated and determined in GIMP [39], before volume was approximated as an ellipsoid (4/3 * π * W^2^ * L).

### Nutritional assays

Briefly, nutritional assays for lipid, glycogen, and trehalose levels were adapted from previous studies [40–42] and combined to allow for technical and biological replication. After relative humidity treatments, additional cohorts were permitted access to water and 10% sucrose solutions *ad libitum* for 24 hours to represent recovery conditions from these treatments. The colony group in this context represents *A. aegypti* that were subjected to only colony conditions and not any additional RH treatment. For quantification, mosquitoes were collected from the same group, placed in a freezer until death (-20°C), added in groups of 4 to STE buffer (2% Na_2_SO_4_), homogenized (Benchmark, BeadBlaster 24), and aliquoted for lipid (100μL), trehalose (150μL), and glycogen (150μL). Six groups in biological triplicates and two standard curves in technical duplicate were distributed across two 96-well plates (Zinsser). Absorbance was determined on a microplate reader (Biotek, Synergy H1) at 525 and 625nm for lipids and carbohydrates respectively. Due to the nested nature of the biological sample replicates, each group was replicated at least thrice on the two-plate design.

### Statistical analyses

Data management was completed in Excel [43] and R [44] through plyr [45], tidyr [46], dplyr [47], and Rmisc [48] packages. Figures were made in R using ggplot2 [49], in Excel [43], and with CirGO, before finalization in GIMP [39] and Inkscape [50]. Tables were made in Excel [43]. R (version 4.0.2) was used to complete appropriate statistical analyses [44].

## Results

### Gene ontology reveals slight differences between midgut groups

Our groups consisted of non-bloodfed (N), and bloodfed (Y) mosquitoes held at either 75% RH (7) or 100% RH (1). Our analyses identified hundreds of genes with differentially expressed transcripts between midgut group comparisons, revealing relatively constrained functionality within the midgut regardless of dehydration or bloodfeeding (Table 2). Despite the three-fold number of genes identified between the dehydrated and non-dehydrated midguts of non-bloodfed *A. aegypti* (237 genes), the comparison between dehydrated and non-dehydrated bloodfed midguts had the lowest number of differentially expressed genes, with less than 80 total genes identified (Table 2). These comparisons underscore the similarities in dehydrated and non-dehydrated midgut functionality within three hours pbf (Table 2).

**Table 2:** Group comparison information regarding significantly expressed genes, Gene Ontology (GO) pathways, and REVIGO terms. Gene lists, GO pathways, and REVIGO terms were generated from transcripts identified by any pipeline. Specific information can be found in Supplementary Tables 1-4. Group N1Y1 represents comparisons between the non-bloodfed/non-dehydrated and the bloodfed/non-dehydrated groups; N7N1, non-bloodfed/dehydrated and non-bloodfed/non-dehydrated groups; Y7Y1, bloodfed/dehydrated and bloodfed/non-dehydrated groups; N7Y7, non-bloodfed/dehydrated and bloodfed/dehydrated groups.

| Group | Comparison | Genes | GO Pathways | REVIGO Terms |
| --- | --- | --- | --- | --- |
| Y1N1 | Y1/N1 | 145 | 7 | 4 |
|  | N1/Y1 | 62 | 3 | 3 |
| N7N1 | N7/N1 | 146 | 4 | 2 |
|  | N1/N7 | 91 | 13 | 5 |
| Y7Y1 | Y7/Y1 | 37 | 0 | 0 |
|  | Y1/Y7 | 40 | 0 | 0 |
| N7Y7 | N7/Y7 | 390 | 8 | 5 |
|  | Y7/N7 | 281 | 29 | 4 |

All comparisons showed GO differences except for the contrasts between Y7 and Y1 groups, indicating that regardless of the level of dehydration status experienced in this study, bloodmeal processing in the midgut was remarkably similar (Figure 1; Supplementary Figure 1). The primary non-bloodfed N7_N1 comparison revealed cell and membrane interactions (Figure 1A), while the N1_N7 comparison showed persistent changes to ion channel activity (Figure 1B). The N1_Y1 comparison showed differences in developmental and regulatory genes (Supplementary Figure 2A), Y1_N1 revealed GO terms consistent with bloodmeal breakdown (Supplementary Figure 2B), N7_Y7 showed changes in protein binding and transcription (Supplementary Figure 2C), and Y7_N7 also uncovered GO terms associated with bloodfeeding as well as a number of terms relating to snRNPs and RNA functionality (Supplementary Figure 2).

**Figure 1:**
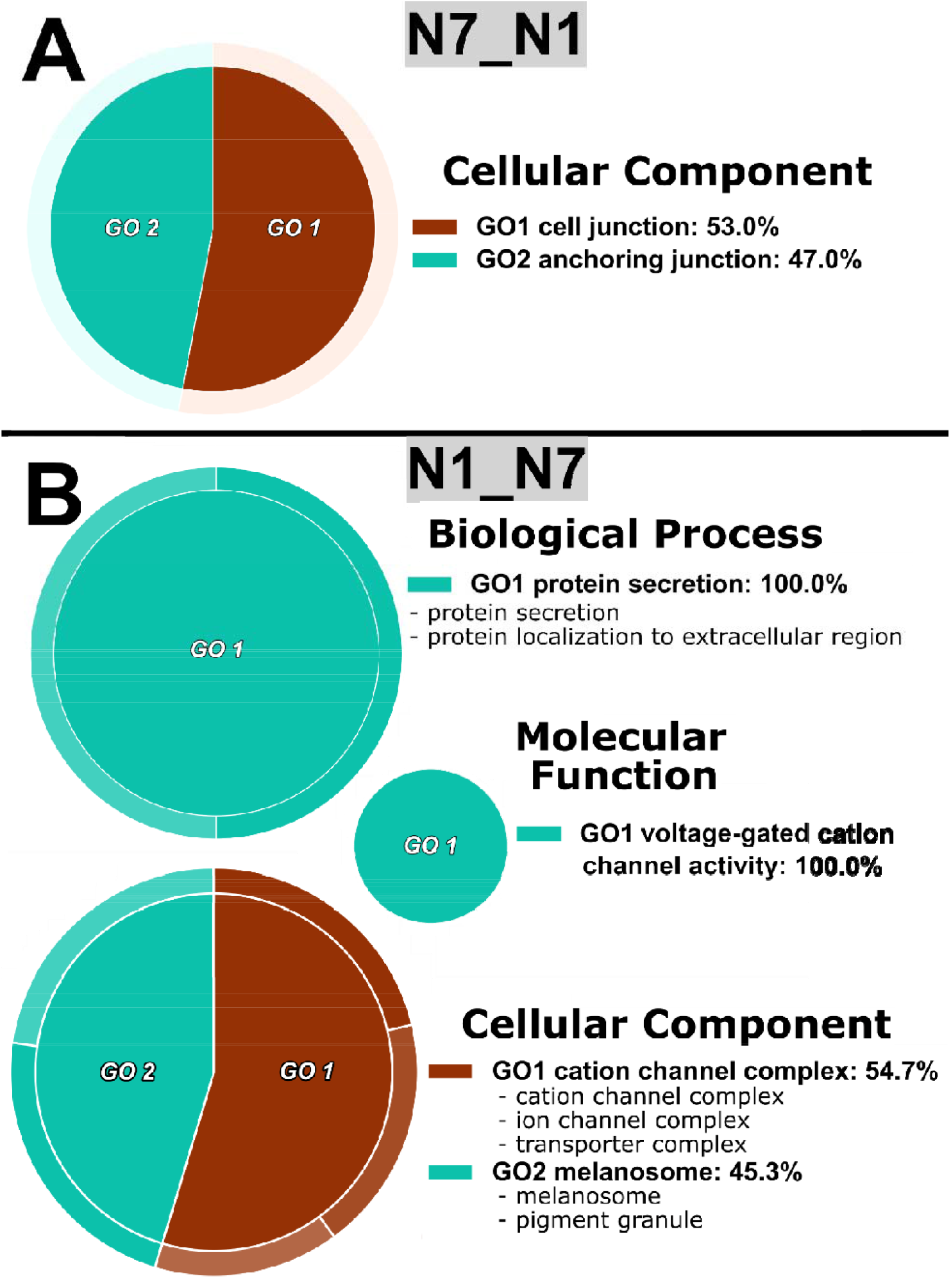
Functional enrichment analyses for non-bloodfed *A. aegypti* midguts. A, circular gene ontology (CirGO) representations of reduced and visualized gene ontology (REVIGO) terms in the non-bloodfed/dehydrated group over the non-bloodfed/non-dehydrated group (N7_N1); B, CirGO-REVIGO representations for the non-bloodfed/non-dehydrated group over the non-bloodfed/dehydrated group over (N1_N7). REVIGO groupings are included in Supplementary Table 3 and significant g:Profiler terms are included in Supplementary Table 4 with “intersections” indicating the genes responsible for GO categorization. CLC labels represent significant transcripts identified with the QIAGEN CLC pipeline; DS, the DESeq2-kallisto pipeline; and DK, the DESeq2-Sailfish pipeline.

In both the dehydrated and non-dehydrated comparisons between bloodfed and non-bloodfed *A. aegypti*, numerous transcripts directly associated with bloodmeal processing (e.g., trypsin, peritrophin, etc.) were upregulated in the bloodfed group, while a limited and lowly expressed set were significantly differentiated in the non-bloodfed group (Supplementary Figure 3). When comparing non-bloodfed groups, dehydrated *A. aegypti* had considerably more, and more highly expressed, transcripts than the non-dehydrated group (Figure 2A). In our dehydrated comparison (Supplementary Figure 3B), the non-bloodfed group also showed considerably more transcripts than the non-bloodfed, non-dehydrated group in a similar comparison (Supplementary Figure 3B; Table 2). The dehydrated group also expressed significant transcripts related to transporters and apoptosis while the non-dehydrated control had lowly-expressed phosphatases with high fold-changes (Figure 2A). When comparing bloodfed groups, there were only a couple dozen differentially expressed genes between the non-dehydrated and dehydrated groups, while all the transcripts had low mean expression values (Figure 2B). Furthermore, the non-dehydrated bloodfed group consisted of transcripts encoding cytoskeletal/structural elements (e.g., rhophilin-2, Lasp, etc.) and the dehydrated bloodfed group featured differential regulation of ion transporters and kinases (Figure 2B). The dehydrated comparison between non-bloodfed and bloodfed *A. aegypti* showed stark similarities to the non-dehydrated bloodfeeding comparison in regard to bloodmeal processing (Supplementary Figure 3).

**Figure 2:**
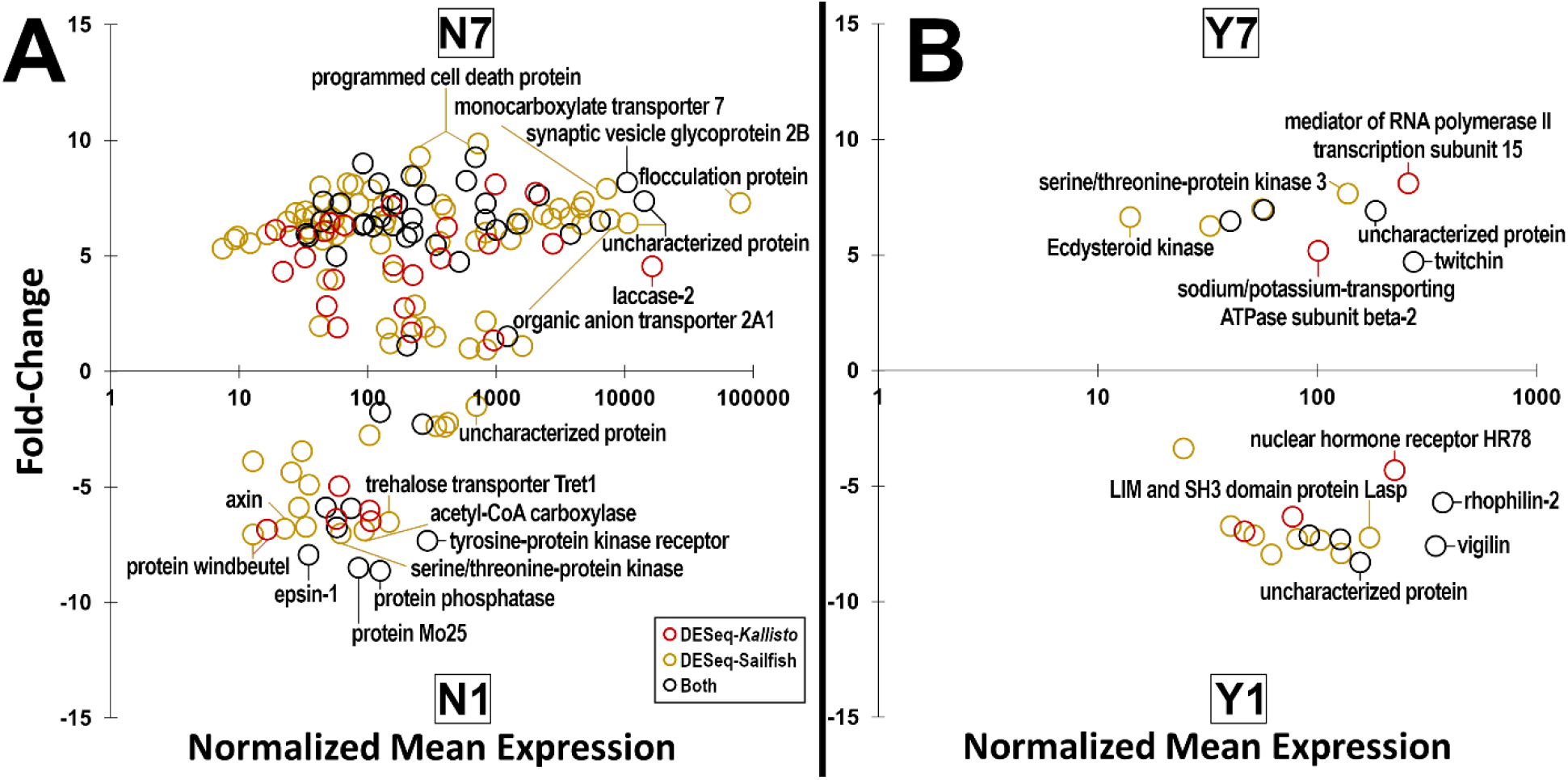
Fold-change and normalized mean expression comparisons for all significantly expressed genes identified by DESeq2 pipelines. A, comparison between the non-bloodfed/dehydrated group over the non-bloodfed/non-dehydrated group (N7_N1); B, comparison between the bloodfed/non-dehydrated group over the bloodfed/dehydrated (Y1_Y7). Yellow circles denote genes that were identified through the DESeq-Sailfish pipeline; red circles, DESeq-*Kallisto* pipeline; and black circles were genes identified by both pipelines, with the highest mean expression pipeline used. Significantly expressed transcripts are included in Supplementary Table 1.

### Post-dehydration bloodfeeding shifts hemolymph osmolality back to control levels

Osmolality in the hemolymph increased as mosquitoes lost water, but within 1h pbf, hemolymph osmolality returned to control levels in dehydrated-then-bloodfed mosquitoes (Figure 3A). No alterations to lipid, glycogen, or the primary hemolymph carbohydrate, trehalose, were identified (Supplementary Figure 4). Finally, no distinguishable volume changes were identified in the dissected midguts of non-dehydrated/control-then-bloodfed, dehydrated-then-bloodfed, nor colony-then-bloodfed mosquitoes (Figure 3B).

**Figure 3:**
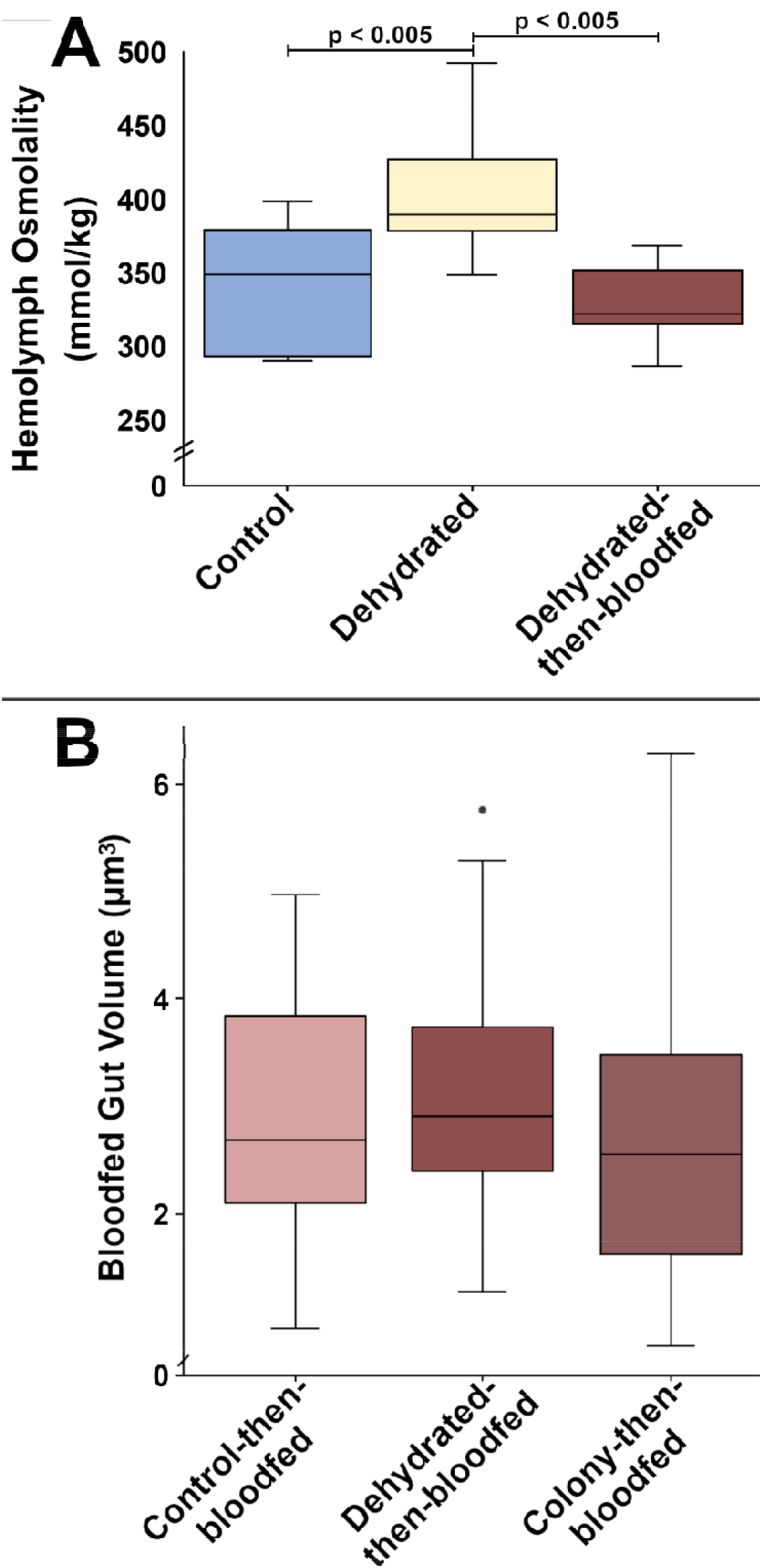
Hemolymph osmolality and bloodfed midgut volume for *A. aegypti* subjected to various treatments. A, hemolymph osmolality for control, dehydrated, and post-dehydration bloodfed *A. aegypti* (N = 30); B, midgut size comparisons for bloodfed *A. aegypti* after 18h of exposure to control, dehydrating, or colony conditions (N = 86). Significance was determined via ANOVA and Tukey’s HSD test.

## Discussion

Through bloodfeeding, mosquitoes have been afforded flexibility to the regulation of nutrients, reproductive output, survival, and more when compared to non-bloodfeeding organisms [51,52]. For example, female mosquitoes with diminished nutritional reserves are capable of diverting nutrients from a bloodmeal to supplement existing levels, but do so at the expense of reproductive output [53]. Likewise, stress related to teneral nutritional reserves may result in differentially utilized nutrients [25]. It is therefore understandable that mosquitoes stressed with acute or persistent dehydration have adapted numerous mechanisms to combat this influence [17]. A recent study investigating the physiological effects of dehydration demonstrated that water loss plays an integral role in mosquito reproduction, survival, water content regulation, and vectorial capacity [7]. In this study, we expand on these findings by exploring the potential underlying mechanisms by which these physiological changes may occur, through investigation of transcriptomic, volumetric, and osmolality changes at the midgut interface.

A previous study on the whole-body transcriptome of non-bloodfed dehydrated *C. pipiens* showed that many significantly upregulated pathways were related to carbohydrate metabolism [1]. These carbohydrate metabolism pathway alterations clearly corroborate the findings in another study showing that repeated bouts of dehydration resulted in reduced levels of stored carbohydrates and lipids in *C. pipiens* [8]. When sugar and water were withheld and mosquitoes were permitted or prohibited to bloodfeed, proteins were consistently altered [7], but our research showed that other micronutrients including trehalose, glycogen, and lipids were no different between groups (Supplementary Figure 4). The lack of significant changes to nutrition were likely the result from the short interval in which the metabolic assays were completed (<18h after experimental onset), but nonetheless represent responses to water loss, not nutritional depletion. Benoit et al., (2010) dehydrated non-bloodfed *C. pipiens* to the point of 25% water loss (comparable water loss to our study) then allowed them to recover before taking nutrient levels and likewise saw no differences in lipids, glycogen, protein, or sugar levels. Although the midgut-specific focus of the sequencing in this research limited the breadth at which carbohydrate metabolism pathways could be discovered, the resolution at which the expressional analyses were performed allowed us to thoroughly investigate the effects of bloodfeeding and dehydration at the intersection of the midgut. Through analysis of the underlying mechanisms, we have facilitated a more thorough understanding on how mosquitoes respond to dehydration stress in the context of 1) water and nutrient utilization and 2) bloodmeal protein utilization. This mechanistic knowledge provides much needed context for recent discoveries involving the effects of dehydration stress on survival, reproduction, and vectorial capacity, within medically-important mosquitoes species [1,7].

To process a bloodmeal, which is composed of 80-87% water and approximately 90% protein composition in the remaining dry mass, mosquitoes must promptly and efficiently regulate these abundant resources [54,55]. Under normal conditions, approximately 40% of water, sodium (Na), and chloride (Cl) derived from a bloodmeal are reportedly excreted within the first two hours pbf [27]. However, as *A. aegypti* become dehydrated, pbf diuresis substantially decreases [7], likely resulting in increased urine retention by the Malpighian tubules. This information coupled with our osmolality findings taken one-hour pbf indicate that *A. aegypti* can exchange ions and extract water from a bloodmeal when necessary to combat dehydration. While ions are actively transferred through the midgut, as indicated by differential expression of ion transporters in this study, water transfer from the more dilute human blood into the hemolymph may occur passively due to osmolality differences [27,56]. The excessive quantities of water and protein in a bloodmeal afford flexibility to mosquitoes, allowing for excretion or rapid replacement of previously lost water. The increased retention of bloodmeal components, as seen in this study through reequilibrated hemolymph osmolality and transcriptional regulation of ion transporters, is also corroborated by previous studies reporting reduced diuresis as well as by high variability observed in the dry masses of dehydrated mosquitoes [this study,1,7]. Specifically, our study shows that numerous genes consistent with ion channel activity were differentially regulated between our non-dehydrated and dehydrated groups and that bloodmeal processing (e.g., trypsin, peritrophin) genes were differentially regulated in our bloodfed groups. Our osmolality data paired with the expression of ion transporters during *A. aegypti* dehydration, further underscores the importance of water content regulation in mosquitoes.

As for protein utilization, a considerable amount of enzymatic/proteolytic activity occurs in the ectoperitrophic space, and very little activity in the blood-filled midgut homogenates [57,58]. A number of these processes are implicated in our transcriptional analyses (e.g., peritrophin, trypsin, etc.). Additional transcripts such as ion transporters and kinases offer insight into the potential means through which *A. aegypti* may compensate for dehydration and bloodfeeding stress at the midgut interface. In our comparison between bloodfed groups, the dehydrated group had increased expression in a number of kinases over the non-dehydrated group. Of particular interest, one specific gene (AAEL012685-RC) encoded an ecdysteroid kinase (the family including ecdysteroid 22-kinase), which closely identifies with juvenile hormone-inducible proteins and hypothetical proteins found across an array of other medically-important mosquito species (e.g., *Anopheles gambiae, Culex pipiens, Aedes, albopictus*, etc.; Supplementary Table 5). This may offer additional insight into the reasons behind reduced egg production observed in dehydrated mosquitoes [7], or potentially into the veiled 20-hydroxyecdysone (20E) signaling pathway. Another over-expressed gene of interest identified in our Y1_Y7 comparison, vigilin (AAEL001421-RA), has been implicated in the formation of RACK1, which is involved in viral RNA binding for DENV genome amplification [59]. Considering the abundance of RNA-involved processes in our Y7_N7 comparison, especially regarding our Y1_N1 comparison, possibilities exist for interactions between imbibed pathogens and the genes expressed within dehydrated mosquitoes. However, more research is needed to address the potential for altered processing of a post-dehydration bloodmeal in the event that an imbibed bloodmeal were to contain pathogens such as Mayaro, Zika, or Dengue (DENV) viruses.

Mosquitoes that underwent dehydration stress were predicted to increase WNV infections as a result of increased biting propensity, while in a similar finding, mosquitoes with reduced nutritional reserves had an increased propensity to orally transmit WNV infection [1,18]. We originally postulated that mosquitoes may compensate for dehydration stress by over-indulging on a bloodmeal, resulting in increased permissibility for imbibed pathogens via induced microperforations [60], but our volumetric analyses determined that the midgut was not overfilled immediately after bloodfeeding. These findings, however, do not exclude the influences of gene regulation on pathogen interactions. It is possible that dehydration may prompt the supplementation of pbf water and nutritional reserves at the expense of reproduction [7,53], and that dehydration may also promote increased instances of refeeding in dehydrated mosquitoes, furthering the potential for additional pathogen exposure for both hosts and vectors. Similar to Armstrong et al. (2020), these dehydration-prompted refeedings may promote microperforations to the midgut, resulting in increased pathogen dissemination. Pathogen dissemination may also be encouraged in dehydrated mosquitoes by expedited passage of bloodmeal components through the midgut barrier via active means such as transporter-facilitated efflux and/or via passive means down a concentration gradient with water from relatively dilute blood to the more concentrated hemolymph. Furthermore, the previously reported reduction to pbf diuresis in dehydrated mosquitoes may continue to alter pathogen interactions within the mosquito via increased bloodmeal retention [7]. To address these possibilities, more research should be completed on the direct influence of dehydration as well as the effects of dehydration-induced refeeding on midgut permissibility to, and downstream retention of, pathogens. Hopefully, these results may be used to continue addressing the gaps in knowledge regarding the impact of dehydration on arthropod-borne disease transmission that still exist. Additional information on the direct interaction between pathogens and dehydrated mosquitoes, especially at the midgut interface, is sorely needed.

## Conclusions

Mosquitoes must meticulously regulate water content to maintain homeostasis, especially after imbibing a bloodmeal. These dynamics become particularly interesting in dehydrating conditions, with a recent study reporting that 70-90% of the largest bloodmeals taken by *A. aegypti* and *C. pipiens* (as indicated by hemoglobin content) were found in dehydrated mosquitoes [7]. However, in this study, we saw no indication of enlargement in dehydrated *A. aegypti* midguts, further indicating the expedited processing of post-dehydration bloodmeals. Taken together with the knowledge that *A. aegypti* are also known to reduce pbf diuresis when dehydrated [7], these results indicate an ability to begin bloodmeal processing for rehydration during or immediately after feeding. This may result in an overall greater intake and retention of a post-dehydration bloodmeal, all while lost water is replenished and maximum midgut size remains unsurpassed. Although *A. aegypti* did not undergo diuresis while feeding as *Anopheles* species do, alterations in GO pathways, underlying genes, bloodmeal processing, and retention in dehydrated *A. aegypti* indicate that similar processes may be involved. Considering the possibility of dehydrated mosquitoes to imbibe and expeditiously process pathogens alongside bloodmeal components, as well as the potential for more direct vector-pathogen interactions, more research on pathogen ingestion and dissemination in this context remains intriguing and necessary.

## Acknowledgements

This research was supported by the National Institute of Allergy and Infectious Diseases of the National Institutes of Health, Award Number R01AI148551. The content reported here is the sole responsibility of the authors and not necessarily a representation of the views of the National Institutes of Health.

## CRediT authorship contribution statement

**Christopher J. Holmes:** Conceptualization, Methodology, Software, Validation, Formal analysis, Investigation, Resources, Data curation, Writing – original draft, Writing – review & editing, Visualization, Supervision, Project administration.

**Elliott S. Brown:** Conceptualization, Methodology, Software, Validation, Formal analysis, Investigation, Data curation.

**Dhriti Sharma:** Conceptualization, Methodology, Investigation, Data curation. **Matthew Warden:** Conceptualization, Methodology, Investigation, Data curation. **Atit Pathak:** Methodology, Investigation, Data curation.

**Blaine Payton:** Investigation, Data curation.

**Quynh Nguyen:** Investigation, Data curation.

**Austin A. Spangler:** Methodology, Investigation, Data curation.

**Jaishna Sivakumar:** Investigation, Data curation.

**Jacob M. Hendershot:** Investigation.

**Joshua B. Benoit:** Conceptualization, Methodology, Validation, Resources, Writing – review & editing, Visualization, Supervision, Project administration, Funding acquisition.

## References

1. Hagan, R.W.; Didion, E.M.; Rosselot, A.E.; Holmes, C.J.; Siler, S.C.; Rosendale, A.J.; Hendershot, J.M.; Elliot, K.S.B.; Jennings, E.C.; Nine, G.A.; et al. Dehydration Prompts Increased Activity and Blood Feeding by Mosquitoes. Sci. Rep. 2018, 8, 1–12, doi:10.1038/s41598-018-24893-z.

2. Canyon, D. V; Hii, J.L.K.; Müller, R. Adaptation of Aedes aegypti (Diptera: Culicidae) Oviposition Behavior in Response to Humidity and Diet. J. Insect Physiol. 1999, 45, 959–964.

3. Costa, E.A.P.D.A.; Santos, E.M.D.M.; Correia, J.C.; Albuquerque, C.M.R. De Impact of Small Variations in Temperature and Humidity on the Reproductive Activity and Survival of Aedes aegypti (Diptera, Culicidae). Rev. Bras. Entomol. 2010, 54, 488–493, doi:10.1590/S0085-56262010000300021.

4. Khan, A.A.; Maibach, H.I. A Study of the Probing Response of Aedes aegypti. 4. Effect of Dry and Moist Heat on Probing. J. Econ. Entomol. 1971, 64, 442–443.

5. Rowley, W.A.; Graham, C.L. The Effect of Temperature and Relative Humidity on the Flight Performance of Female Aedes aegypti. J. Insect Physiol. 1968, 14, 1251–1257, doi:10.1016/0022-1910(68)90018-8.

6. Parker, A.H. The Effect of a Difference in Temperature and Humidity on Certain Reactions of Female Aedes aegypti (L.). Bull. Entomol. Res. 1952, 43, 221–229, doi:10.1017/S0007485300030698.

7. Holmes, C.J.; Brown, E.S.; Sharma, D.; Nguyen, Q.; Spangler, A.A.; Pathak, A.; Payton, B.; Warden, M.; Shah, A.J.; Shaw, S.; et al. Bloodmeal Regulation in Mosquitoes Curtails Dehydration-Induced Mortality, Altering Vectorial Capacity. J. Insect Physiol. 2022, 137, 104363, doi:10.1016/j.jinsphys.2022.104363.

8. Benoit, J.B.; Patrick, K.R.; Desai, K.; Hardesty, J.J.; Krause, T.B.; Denlinger, D.L. Repeated Bouts of Dehydration Deplete Nutrient Reserves and Reduce Egg Production in the Mosquito Culex pipiens. J. Exp. Biol. 2010, 213, 2763–2769, doi:10.1242/jeb.044883.

9. Reidenbach, K.R.; Cheng, C.; Liu, F.; Liu, C.; Besansky, N.J.; Syed, Z. Cuticular Differences Associated with Aridity Acclimation in African Malaria Vectors Carrying Alternative Arrangements of Inversion 2La. Parasites and Vectors 2014, 7, 1–13, doi:10.1186/1756-3305-7-176.

10. Canyon, D. V.; Muller, R.; Hii, J.L.K. Aedes aegypti Disregard Humidity-Related Conditions with Adequate Nutrition. Trop. Biomed. 2013, 30, 1–8.

11. Dow, R.P.; Gerrish, G.M. Day-to-Day Change in Relative Humidity and the Activity of Culex nigripalpus (Diptera: Culicidae). Ann. Entomol. Soc. Am. 1970, 63, 995–999, doi:10.1093/aesa/63.4.995.

12. Lyons, C.L.; Coetzee, M.; Terblanche, J.S.; Chown, S.L. Desiccation Tolerance as a Function of Age, Sex, Humidity and Temperature in Adults of the African Malaria Vectors Anopheles arabiensis and Anopheles funestus. J. Exp. Biol. 2014, 217, 3823–3833, doi:10.1242/jeb.104638.

13. Kumar, M. Effect of Temperature and Humidity on Life Cycle Duration of Culex quinquefasciatus Say (DipteraL: Culicidae) at Muzaffarpur (Bihar), India. Adv. Biores. 2015, 6, 103–105, doi:10.15515/abr.0976-4585.6.6.103105.

14. Leeson, H.S. Longevity of Anopheles maculipennis Race Atroparvus, van Thiel, at Controlled Temperature and Humidity after One Blood Meal. Bull. Entomol. Res. 1939, 30, 295–301.

15. Mayne, B. Notes on the Influence of Temperature and Humidity on Oviposition and Early Life of Anopheles. Public Health Rep. 1926, 41, 986–990.

16. Ruiz, M.O.; Chaves, L.F.; Hamer, G.L.; Sun, T.; Brown, W.M.; Walker, E.D.; Haramis, L.; Goldberg, T.L.; Kitron, U.D. Local Impact of Temperature and Precipitation on West Nile Virus Infection in Culex Species Mosquitoes in Northeast Illinois, USA. Parasites and Vectors 2010, 3, 1–16, doi:10.1186/1756-3305-3-19.

17. Holmes, C.J.; Benoit, J.B. Biological Adaptations Associated with Dehydration in Mosquitoes. Insects 2019, 10, 375, doi:10.3390/insects10110375.

18. Vaidyanathan, R.; Fleisher, A.E.; Minnick, S.L.; Simmons, K.A.; Scott, T.W. Nutritional Stress Affects Mosquito Survival and Vector Competence for West Nile Virus. VectorBorne Zoonotic Dis. 2008, 8, 727–732, doi:10.1089/vbz.2007.0189.

19. Briegel, H.; Hörler, E. Multiple Blood Meals as a Reproductive Strategy in Anopheles (Diptera: Culicidae). J. Med. Entomol. 1993, 30, 975–985, doi:10.1093/jmedent/30.6.975.

20. Lea, A.O.; Briegel, H.; Lea, H.M. Arrest, Resorption, or Maturation of Oocytes in Aedes aegypti: Dependence on the Quantity of Blood and the Internval between Blood Meals. Physiol. Entomol. 1978, 3, 309–316.

21. Paz, S. Climate Change Impacts on West Nile Virus Transmission in a Global Context. Philos. Trans. R. Soc. B Biol. Sci. 2015, 370, 1–11, doi:10.1098/rstb.2013.0561.

22. Dodson, B.L.; Rasgon, J.L. Vector Competence of Anopheles and Culex Mosquitoes for Zika Virus. PeerJ 2017, 5, e3096, doi:10.7717/peerj.3096.

23. Tingström, O.; Wesula Lwande, O.; Näslund, J.; Spyckerelle, I.; Engdahl, C.; Von Schoenberg, P.; Ahlm, C.; Evander, M.; Bucht, G. Detection of Sindbis and Inkoo Virus RNA in Genetically Typed Mosquito Larvae Sampled in Northern Sweden. Vector-Borne Zoonotic Dis. 2016, 16, 461–467, doi:10.1089/vbz.2016.1940.

24. Liu, K.; Dong, Y.; Huang, Y.; Rasgon, J.L.; Agre, P. Impact of Trehalose Transporter Knockdown on Anopheles gambiae Stress Adaptation and Susceptibility to Plasmodium falciparum Infection. Proc. Natl. Acad. Sci. 2013, 110, 17504–17509, doi:10.1073/pnas.1316709110.

25. Naksathit, A.T.; Edman, J.D.; Scott, T.W. Utilization of Human Blood and Sugar as Nutrients by Female Aedes aegypti (Diptera: Culicidae). J. Med. Entomol 1999, 36, 13–17.

26. Drake, L.L.; Boudko, D.Y.; Marinotti, O.; Carpenter, V.K.; Dawe, A.L.; Hansen, I.A. The Aquaporin Gene Family of the Yellow Fever Mosquito, Aedes aegypti. PLoS One 2010, 5, 1–9, doi:10.1371/journal.pone.0015578.

27. Williams, J.C.; Hagedorn, H.H.; Beyenbach, K.W. Dynamic Changes in Flow Rate and Composition of Urine during the Post-Bloodmeal Diuresis in Aedes aegypti (L.). J. Comp. Physiol. B 1983, 153, 257–265, doi:10.1007/BF00689629.

28. Matthews, B.J.; Dudchenko, O.; Kingan, S.B.; Koren, S.; Antoshechkin, I.; Crawford, J.E.; Glassford, W.J.; Herre, M.; Redmond, S.N.; Rose, N.H.; et al. Improved Reference Genome of Aedes aegypti Informs Arbovirus Vector Control. Nature 2018, 563, 501–507, doi:10.1038/s41586-018-0692-z.

29. Afgan, E.; Baker, D.; Batut, B.; Van Den Beek, M.; Bouvier, D.; Ech, M.; Chilton, J.; Clements, D.; Coraor, N.; Grüning, B.A.; et al. The Galaxy Platform for Accessible, Reproducible and Collaborative Biomedical Analyses: 2018 Update. Nucleic Acids Res. 2018, 46, W537–W544, doi:10.1093/nar/gky379.

30. Andrews, S. FastQC: A Quality Control Tool for High Throughput 2015.

31. Bolger, A.M.; Lohse, M.; Usadel, B. Trimmomatic: A Flexible Trimmer for Illumina Sequence Data. Bioinformatics 2014, 30, 2114–2120, doi:10.1093/bioinformatics/btu170.

32. Bray, N.L.; Pimentel, H.; Melsted, P.; Pachter, L. Near-Optimal Probabilistic RNA-Seq Quantification. Nat. Biotechnol. 2016, 34, 525–527, doi:10.1038/nbt.3519.

33. Patro, R.; Mount, S.M.; Kingsford, C. Seq Reads Using Lightweight Algorithms. Nat. Biotechnol. 2014, 32, 462–464, doi:10.1038/nbt.2862.Sailfish.

34. Love, M.I.; Huber, W.; Anders, S. Moderated Estimation of Fold Change and Dispersion for RNA-Seq Data with DESeq2. Genome Biol. 2014, 15, 1–21, doi:10.1186/s13059-014-0550-8.

35. Reimand, J.; Kull, M.; Peterson, H.; Hansen, J.; Vilo, J. G:Profiler-a Web-Based Toolset for Functional Profiling of Gene Lists from Large-Scale Experiments. Nucleic Acids Res. 2007, 35, 193–200, doi:10.1093/nar/gkm226.

36. Supek, F.; Bošnjak, M.; Škunca, N.; Šmuc, T. Revigo Summarizes and Visualizes Long Lists of Gene Ontology Terms. PLoS One 2011, 6, doi:10.1371/journal.pone.0021800.

37. Kuznetsova, I.; Lugmayr, A.; Siira, S.J.; Rackham, O.; Filipovska, A. CirGO: An Alternative Circular Way of Visualising Gene Ontology Terms. BMC Bioinformatics 2019, 20, 1–7, doi:10.1186/s12859-019-2671-2.

38. Detinova, T.S.; Beklemishev, W.N.; Bertram, D.S. Age-Grouping Methods in Diptera of Medical Importance With Special Reference to Some Vectors of Malaria. World Heal. Organ. Monogr. Ser. 1962, 47, 1–213.

39. The GIMP Development Team GIMP.

40. Rivers, D.B.; Denlinger, D.L. Redirection of Metabolism in the Flesh Fly, Sarcophaga bullata, Following Envenomation by the Ectoparasitoid Nasonia vitripennis and Correlation of Metabolic Effects with the Diapause Status of the Host. J. Insect Physiol. 1994, 40, 207–215, doi:10.1016/0022-1910(94)90044-2.

41. Van Handel, E. Rapid Determination of Glycogen and Sugars in Mosquitoes. J. Am. Mosq. Control Assoc. 1985, 1, 299–301.

42. Van Handel, E. Rapid Determination of Total Lipids in Mosquitoes. J. Am. Mosq. Control Assoc. 1985, 1, 302–304.

43. Microsoft Corporation Microsoft Excel.

44. R Core Team R: A Language and Environment for Statistical Computing 2021.

45. Wickham, H. The Split-Apply-Combine Strategy for Data Analysis. J. Stat. Softw. 2011, 40, 1–29.

46. Wickham, H. Tidyr: Tidy Messy Data 2020.

47. Wickham, H.; Francois, R.; Henry, L.; Müller, K. Dplyr: A Grammar of Data Manipulation 2017.

48. Hope, R.M. Rmisc: Rmisc: Ryan Miscellaneous 2013.

49. Wickham, H. Ggplot2: Elegant Graphics for Data Analysis; Springer-Verlag New York, 2009; ISBN 978-0-387-98140-6.

50. Inkscape Project Inkscape 2020.

51. Nayar, J.K.; Sauerman, D.M. The Effects of Nutrition on Survival and Fecundity on Florida Mosquitoes. J. Med. Ent 1975, 12, 99–103.

52. Holt, R.A.; Subramanian, G.M.; Halpern, A.; Sutton, G.G.; Charlab, R.; Nusskern, D.R.; Wincker, P.; Clark, A.G.; Ribeiro, M.C.; Wides, R.; et al. The Genome Sequence of the Malaria Mosquito Anopheles gambiae. October 2002, 298.

53. Foster, W.A. Mosquito Sugar Feeding and Reproductive Energetics. Annu. Rev. Entomol. 1995, doi:10.1146/annurev.ento.40.1.443.

54. Lehane, M.J. The Biology of Blood-Sucking in Insects, Second Edition; Cambridge University Press: New York, 2005; ISBN 9780511610493.

55. Sanders, H.R.; Foy, B.D.; Evans, A.M.; Ross, L.S.; Beaty, B.J.; Olson, K.E.; Gill, S.S. Sindbis Virus Induces Transport Processes and Alters Expression of Innate Immunity Pathway Genes in the Midgut of the Disease Vector, Aedes aegypti. Insect Biochem. Mol. Biol. 2005, 35, 1293–1307, doi:10.1016/j.ibmb.2005.07.006.

56. Piermarini, P.M. Renal Excretory Processes in Mosquitoes; 1st ed.; Elsevier Ltd., 2016; Vol. 51; ISBN 9780128024577.

57. Van Handel, E.; Romoser, W.S. Proteolytic Activity in the Ectoperitrophic Fluid of Blood-fed Culex nigripalpus. Med. Vet. Entomol. 1987, 1, 251–255, doi:10.1111/j.1365-2915.1987.tb00351.x.

58. Clements, A.N. The Biology of Mosquitoes: Development, Nutrition and Reproduction; Chapman & Hall: London, 1992; Vol. 1;.

59. Brugier, A.; Hafirrassou, M.L.; Pourcelot, M.; Baldaccini, M.; Kril, V.; Couture, L.; Kümmerer, B.M.; Gallois-Montbrun, S.; Bonnet-Madin, L.; Vidalain, P.-O.; et al. RACK1 Associates with RNA-Binding Proteins Vigilin and SERBP1 to Facilitate Dengue Virus Replication. J. Virol. 2022, 96, doi:10.1128/jvi.01962-21.

60. Armstrong, P.M.; Ehrlich, H.Y.; Magalhaes, T.; Miller, M.R.; Conway, P.J.; Bransfield, A.; Misencik, M.J.; Gloria-Soria, A.; Warren, J.L.; Andreadis, T.G.; et al. Successive Blood Meals Enhance Virus Dissemination within Mosquitoes and Increase Transmission Potential. Nat. Microbiol. 2020, 5, 239–247, doi:10.1038/s41564-019-0619-y.

