## Supplemental Figures for "Dehydration alters transcript levels in the mosquito midgut, likely facilitating rapid rehydration"

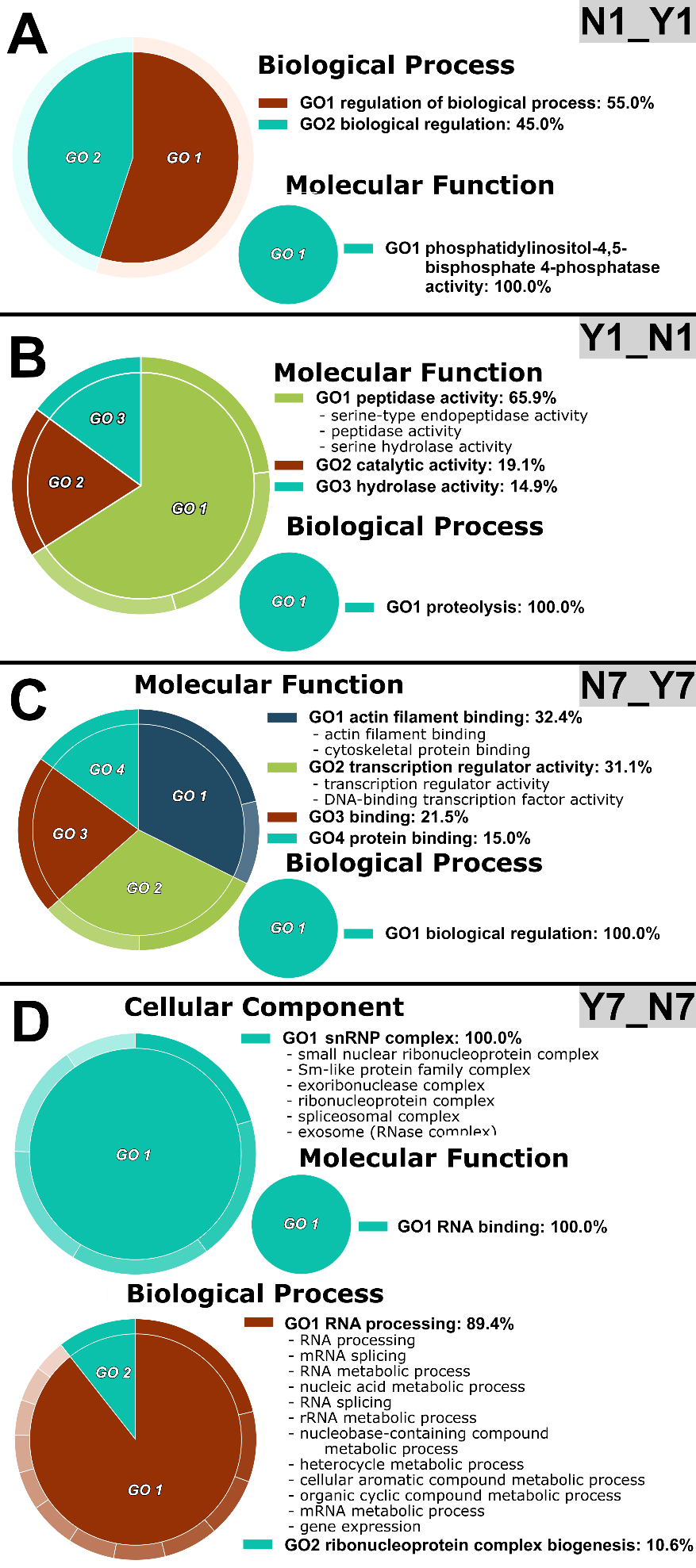

**Supplementary Figure 1:** Functional enrichment analyses. A, CirGO-REVIGO representations for the non-bloodfed/non-dehydrated group over the bloodfed/non-dehydrated group (N1_Y1); B, CirGO-REVIGO representations for the bloodfed/non-dehydrated group over the non-bloodfed/non-dehydrated group (N1_Y1); C, CirGO-REVIGO representations for the non-bloodfed/dehydrated group over the bloodfed/dehydrated group (N7_Y7); D, CirGO-REVIGO representations for the bloodfed/dehydrated group over the non-bloodfed/dehydrated group (Y7_N7). REVIGO groupings are included in Supplementary Table 3 and significant g:Profiler terms are included in Supplementary Table 4 with “intersections” indicating the genes responsible for GO categorization. CLC labels represent significant transcripts identified with the QIAGEN CLC pipeline; DS, the DESeq2-kallisto pipeline; and DK, the DESeq2-Sailfish pipeline.

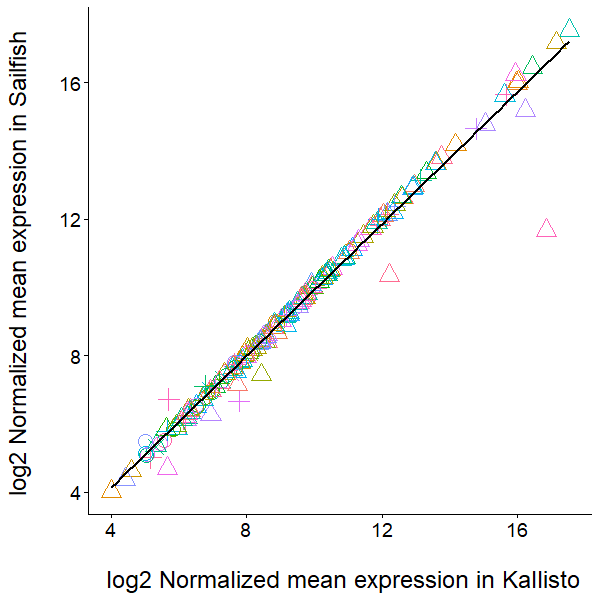

**Supplementary Figure 2:** DESeq2 pipeline comparisons show considerable correlation in shared transcript expression. The Y1_N1 comparison is denoted by circles; Y1_Y7, triangles; N7_N1, squares; N7_Y7, crosses.

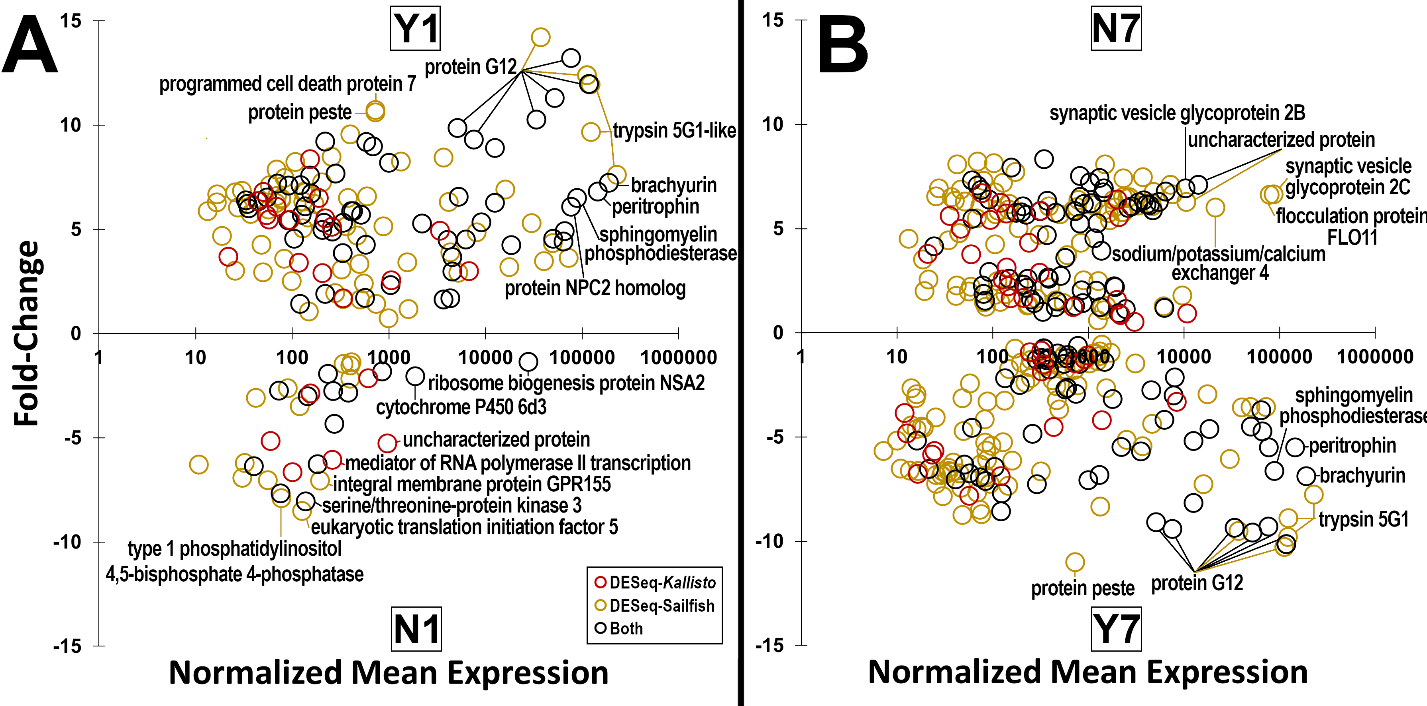

**Supplementary Figure 3:** Fold-change and normalized mean expression comparisons for all significantly expressed genes identified by DESeq2 pipelines. A, comparison between the bloodfed/non-dehydrated group over the non-bloodfed/non-dehydrated group (Y1_N1); B, comparison between the non-bloodfed/dehydrated group over the bloodfed/dehydrated group (N7_Y7). Yellow circles denote genes that were identified through the DESeq-Sailfish pipeline; red circles, DESeq-*Kallisto* pipeline; and black circles were genes identified by both pipelines, with the highest mean expression pipeline used. Significantly expressed transcripts are included in Supplementary Table 1.

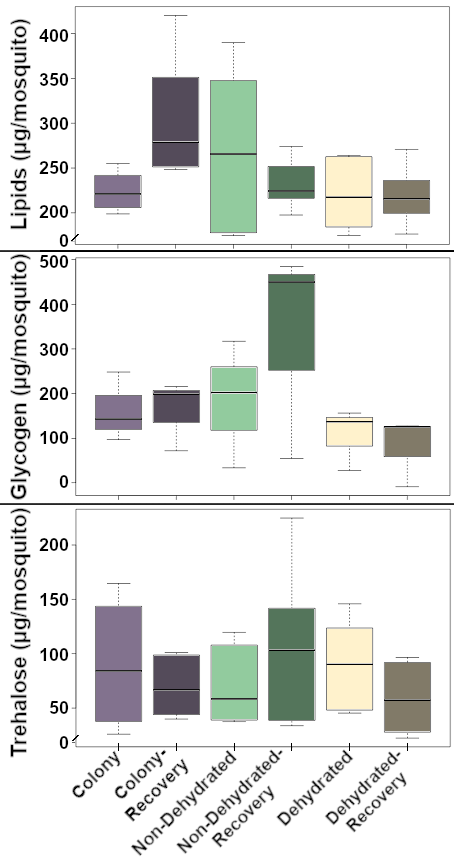

**Supplementary Figure 4:** Macronutrient levels for *A. aegypti* subjected to various treatments.

**Supplementary Table 1:** All significant comparisons (p-value < 0.01, fold-change >2) between groups. Including gene transcripts, nucleotide ID, protein IDs, and relevant protein description. In each Excel worksheet, the initial term (e.g., Y1) represents increased expression over the second term (e.g., N1) in a sequence (e.g., Y1_N1 represents Y1 gene expression over N1 expression). CLC, abbreviation for genes identified by the CLC pipeline; DK, DESeq2-Kallisto; DS, DESeq2-Sailfish; FC, fold-change; -p, p-value.

**Supplementary Table 2:** All significant DESeq2-identified transcript comparisons (p-value < 0.01) between groups. Including gene transcripts, highest mean expression, fold-change, protein ID, and relevant protein description. Each Excel worksheet represents a group comparison. Red fill represents genes identified by the DESeq2-Kallisto pipeline; yellow, DESeq2-Sailfish; black, both DESeq2-pipelines.

**Supplementary Table 3:** All significant REVIGO representatives for GO terms between groups. Including GO term IDs, GO ID names, and REVIGO-assigned representative category. In each Excel worksheet, the initial term (e.g., Y1) represents increased expression over the second term (e.g., N1) in a sequence (e.g., Y1N1 represents Y1 gene expression over N1 expression), the following term (e.g., BP) denotes the GO abbreviation. GO, abbreviation for gene ontology; BP, biological process; CC, cellular component; MF, molecular function.

**Supplementary Table 4:** All significant GO terms and KEGG pathways identified by g:Profiler between groups. Including term names, term IDs, p-value, number of transcript intersections, and identified intersecting gene transcripts. In each Excel worksheet, the initial term (e.g., Y1) represents increased expression over the second term (e.g., N1) in a sequence (e.g., Y1N1 represents Y1 gene expression over N1 expression), the following term (e.g., BP) denotes the GO abbreviation. GO, abbreviation for gene ontology; BP, biological process; CC, cellular component; MF, molecular function; KEGG, Kyoto Encyclopedia of Genes and Genomes pathways.

**Supplementary Table 5:** BLAST information of proteins associated with an identified ecdysteroid kinase protein (gene ID AAEL012685-RC).
